# New permanent stem cell niche for development and regeneration in a chordate

**DOI:** 10.1101/2023.05.15.540819

**Authors:** Virginia Vanni, Federico Caicci, Anna Peronato, Graziano Martello, Davide Asnicar, Fabio Gasparini, Loriano Ballarin, Lucia Manni

## Abstract

Stem cell niches are defined as the microenvironments where stem cells home, receiving stimuli defining their fate. In vertebrates, stem cell niches are stable and physically confined compartments. *Botryllus schlosseri* is an invertebrate colonial chordate where temporary stem cell niches have been identified in adult individuals that are cyclically resorbed and replaced by a new generation of clonal zooids. *B. schlosseri* also displays remarkable regenerative abilities, being capable of whole-body regeneration, but the cellular source of these processes is still unknown. Here we identified by means of a high-resolution morphological characterization a new putative stem cell niche in the ampullae of the circulatory system acting as a stem cell source during asexual reproduction. Stem cells of the ampullae travel via the circulatory system and contribute to the development of several organs and could explain where stem cells contributing to whole-body regeneration are stored. The ampullae niches are stable during the life cycle and regeneration of *B. schlosseri*, while additional niches of the zooid are dynamically established and colonised by circulating stem cells. Our results reveal an unprecedented dynamicity of stem cell niches in highly regenerative invertebrates.

## Introduction

Colonial tunicates are the only chordates capable of whole-body regeneration, meaning that they can recreate complete individuals from adult stem cells (SCs) [1–5]. Understanding the causes and mechanisms of such powerful regenerative abilities is of interest from an evolutionary perspective (tunicates are the sister group of vertebrates [6–8]), and for potential advances in human regenerative medicine.

*Botryllus schlosseri* (Fig. 1a–b, Supplementary Fig. 1a–b) is a widespread colonial tunicate, displaying such high regenerative abilities [2, 9, 10]. Colonies host three generations of individuals called zooids: filter feeding adults, bearing palleal primary buds which, in turn, hold secondary buds or budlets, the youngest generation of zooids (Fig. 1b, Supplementary Fig. 1a–c). During its constitutive asexual (or blastogenetic) cycle, adult individuals are cyclically resorbed during a process called takeover, and replaced by primary buds, which open their syphons becoming adult individuals. In the meantime, secondary buds become primary buds producing a new generation of buds, guaranteeing in this way constant renewals of zooids [11, 12] (Supplementary Fig. 1c). Whole body regeneration in *B. schlosseri* is triggered when all the zooids of the colony are removed, leaving only the colonial circulatory system (CCS) containing blood cells, the hemocytes [2, 10]. The CCS is composed by blind structures called ampullae and by the marginal vessel that runs at the periphery of the colony, connecting all the individuals through radial vessels. The continuous development of new individuals, and whole-body regeneration must be associated with a high demand of SCs. However, SC niches have been described only in adult zooids [13–15], meaning that they are transitory structures, whose fate is to be resorbed together with the adult individuals that bear them. Moreover, when whole-body regeneration occurs, the SC niches identified until now are absent. Indeed, the source of SCs guiding this regeneration process is still unknown, and a permanent SC niche that would support whole body regeneration has not been yet identified. We therefore hypothesised that such permanent niches could be hosted in the CCS.

**Fig. 1.**
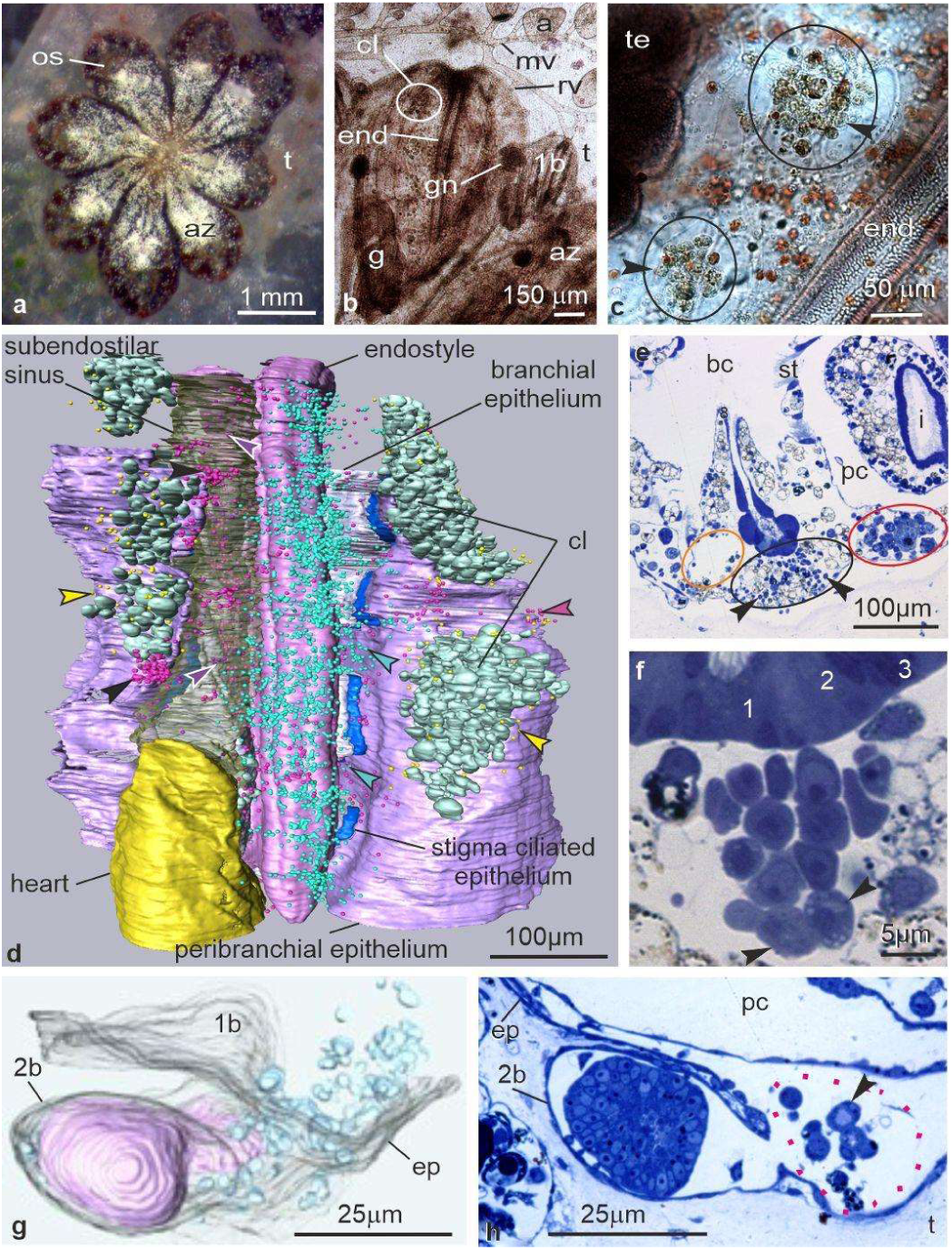
EN, CI and GN *in vivo* and in 3D reconstruction. **a.** Colony of *B. schlosseri*. Dorsal view. az: adult zooid; os: oral syphon; t; tunic. **b-c.** Fixed, whole mount colony in ventral view showing the endostyle (end) and associated niche and cell islands (cI, circles) in an adult zooid (az), and gonad niche (gn) in a primary bud (1b). Arrowheads: phagocytes in cell islands. a: ampulla; g: gut; mv: marginal vessel; rv: radial vessel; t: tunic; te: testis in adult zooid. **d.** 3D reconstruction of the posterior half of the EN and the surrounding tissues of an adult zooid (colony phase 9/8/3). Ventral view. Epidermis omitted to show the EN niche rich in cSCs (blue spots, blue arrowheads). Subendostylar sinus transparent to show a few cSCs in its lumen (purple spots, purple arrowheads). Some cSCs are in the connective tissue (fuchsia spots, fuchsia arrowheads), sometimes aggregated in small nodules (black arrowheads). A few cSCs are in CI (yellow spots, yellow arrowheads). **e.** Transverse histological section of the adult zooid shown in (a). Many cSCs (arrowheads) are in the EN (black circle) whereas the subendostylar sinus (orange circle) does not exhibit them. Red circle: CI. bc: branchial chamber; i: intestine; pc: peribranchial chamber; st: stigma. Toluidine blue. **f.** Rows of cSCs in the EN, with more differentiated ones (arrowheads) far from endostyle. 1-3: endostyle zones. Toluidine blue. **g-h**. GN. 3D reconstruction (**g**) of a secondary bud (2b, stage 3). Anterior view. The inner vesicle (light pink) is connected to the parental peribranchial epithelium. Epidermis (ep) is transparent allowing germ cells (light blue) recognition in the peduncle connecting the bud to the parental primary bud (1b). **h**: transverse histological section of the secondary bud (2b) shown in (g). A previtellogenetic oocytes (arrowhead), surrounded by a layer of primary follicle cells, is in the sinus of the peduncle (dotted red line). Toluidine blue. pc: 1b peribranchial chamber; t: tunic.

Candidate SCs (cSCs) in *B. schlosseri* are recognized on a morphological basis: they are small (among 5-8 µm in diameter), round cells with a high nucleus/cytoplasm ratio, a thin area of cytoplasm without specialisations, such as vacuoles or granules, but presenting few endoplasmic reticulum cisterns and mitochondria. The SC niches identified in the adult zooids of this species are three: the endostyle niche [13] (EN), the cell islands [14] (CIs), and the gonadal niche [15] (GN) (Fig. 1b–c, Supplementary Fig. 1a). Transcriptomic data coupled with transplantation experiments supported the presence of somatic cSCs including hematopoietic cSCs in the EN [13, 16], and proliferation experiments have shown that cSCs coming from these locations can divide and differentiate [13, 14]. However, these niches have never been morphologically characterised, and the actual presence of resident cells together with the morphology of cSCs which proliferate and differentiate within these structures has never been reported. Importantly, this information would confirm the niche nature of these compartments and the possible structural similarity between SC niches of tunicates and vertebrates, as confined areas where cells reside, proliferate and receive the stimuli that orchestrate their fate.

In the current study, we performed a high-resolution analysis through a 3D reconstruction of a CCS fragment and a portion of an adult individual. We characterised for the first time the structure of known SC niches, but we also succeeded in characterising the presence of cSCs in the ampullae of the CCS, which express the orthologous of pluripotent stem cell markers in vertebrates (*cMyc*, *Oct4* and *Sox2*). Therefore, we developed a new method to analyse *in vivo* the contribution of cells from the CCS to the asexual development of *B. schlosseri.* Thanks to this novel cell labelling approach we were able to find a population of cSCs that participate in the development of several tissues and organs. Using an *in vivo* longitudinal approach to follow the fate of sorted (through Fluorescence Activated Cell Sorting, FACS) and transplanted cSCs, we demonstrated that such cSCs home in ampullae and differentiate in tissues of developing buds. Finally, through functional experiments, we proved that ampullae are necessary to sustain the development of buds in absence of adult individuals. Such results support that ampullae are the first permanent SC niche found in *B. schlosseri*.

## The connective tissue ventral to the endostyle is the niche in which cSCs reside, proliferate and differentiate

To identify the localization of cSCs in *B. schlosseri* we performed a novel high-resolution analysis of an entire colony in mid-cycle by means of serial sections, confocal microscopy analysis, and 3D reconstruction (Fig. 1d–f, Supplementary Fig. 2).We studied the EN anatomy, proposed to be located in the anterior endostyle region and in the sinus ventral to it [13, 16]. Cells showing the morphology of cSCs were labelled in the 3D reconstruction. In the sample, cSCs were rarely recognized in the connective tissues of the body wall, but around 2400 cSCs were found in the connective tissue ventral to the reconstructed endostyle, in correspondence of its medial zones (zones 1-3 [17]) (Fig. 1e–f). Many cells were also in the connective tissue lateral to the endostyle. Our results indicate that cSCs of the EN are found all along the antero-posterior axis of the endostyle. We checked the exact location of cSCs with respect to the subendostylar sinus to verify if it was part of the niche, as previously reported [13, 16]. The sinus is located in the ventral body wall, close to the endostyle, on its right, anterior to the heart from which it receives the hemolymph [18]. It is not limited by an endothelium, but by connective tissue that also surrounds the endostyle. Within the sinus we found only 57 cells with a cSCs morphology (out of 2457 total recognised cSCs; purple spots in Fig. 2A), which were probably circulating at the moment of sample fixation. Most of the cSCs (1756 out of 2457; blue spots in Fig. 1d) were in the connective tissue ventral to the endostyle, while other cSCs (644 cSCs; fuchsia spots in Fig. 1d) were found in the connective tissue of the body wall located between the endostyle and CI, but not in body wall sinuses; some of them were grouped in small nodules. In live-imaging movies (Supplementary Video 1), cSCs ventral to the endostyle were seen immobile; few cells moved slowly with respect to cells in the near subendostylar sinus (that vice versa moved quickly in the hemolymph flow), concluding that they were residing, rather than circulating, cells.

**Fig. 2.**
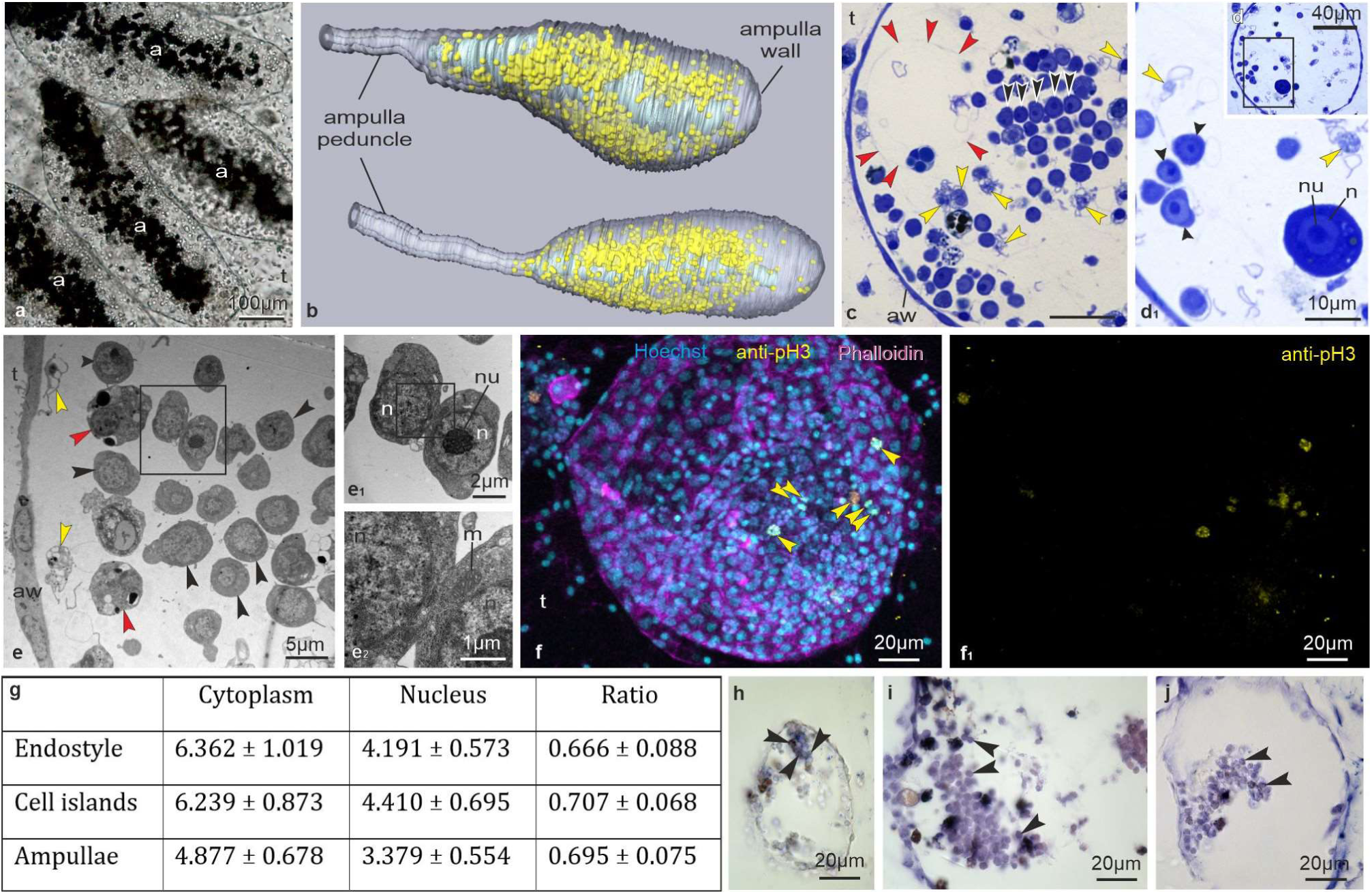
AN organisation and cell proliferation. **a**. *In vivo* ampullae (a). Black spots are immobile pigmented cells, usually found in the connective tissue surrounding the ampulla sinus (hidden by pigmented cells themselves). Note that numerous small cells are recognizable within the ampullae. **b.** 3D reconstruction of two ampullae and part of the peduncle connecting them to the remaining vasculature, obtained by analysing 584 serial sections, 1 µm thick. The ampulla wall is transparent to show the cSCs (yellow spots). The upper and lower ampulla are the n.9 and n.5 in Supplementary Fig. 3a, respectively. **c.** Numerous cSCs are in the connective tissue, together with hyaline amebocytes (yellow arrowheads). A series of cSCs (black arrowheads) form a row of contacting cells. Red arrowheads: ampulla sinus. Transverse section, Toluidine blue. **d-d_1_.** An ampulla (d) containing a previtellogenic oocyte, enlarged in d1. Black arrowheads: cSCs; yellow arrowheads: hyaline hamebocyte. Toluidine blue. **e-e_2_.** Detail of an AN showing a cluster of cSCs (black arrowheads). Yellow arrowheads: hyaline amebocytes; red arrowheads: differentiating morula cells. Two contacting cSCs are enlarged in e_1_. Note the narrow intercellular cleft (enlarged in e**_2_**) at the level of plasmalemma apposition. TEM. m: mitochondrium; n: nucleus; nu: nucleolus. **f-f_1_.** Mitotic cells (arrowheads) in an AN are recognized by the anti-pH3. **g.** Cytoplasm and nucleus diameters (in μm) and nucleus:cytoplasm ratio presented as mean ± standard deviation of cSCs in cell islands, ampullae and endostyle niche. There are no significant differences in the nucleus:cytoplasm ratio of cSCs in the three niches (one-way ANOVA F_(2,50)_ = 1.340, p-value = 0.271). **h-j.** *In-situ* hybridization showing positive cSCs (labelled in blue, black arrowheads) for the expression of *Myc* (j), *Pou3* (k) and *SoxB1* (l) in ampullae.

The collection of serial sections used for the 3D reconstruction allowed us to check the cytological features of all the cSCs located in the reconstructed part of the EN, giving us a snapshot of the niche in a mid-cycle individual (Fig. 1e–f; Supplementary Fig. 2c–h). cSCs were often found in rows in which a gradient of differentiation can be appreciated: cells closest to the endostyle were typical cSCs, whereas those farthest from it exhibited granules inside their cytoplasm. The differentiating cells showed features of young immunocytes and phagocytes, supporting the hematopoietic nature of this niche [16]. At transmission electron microscopy (TEM), we could observe small junctional contacts between cSCs in the form of small areas of juxtaposition, suggesting an active interaction and crosstalk between them (Supplementary Fig. 2e–f). Some formed pairs of identical cells, with large portions of their plasmalemma in strict apposition, as signs of just-completed mitosis (Supplementary Fig. 2g). Other cSCs were in mitosis (cells without recognizable nuclear envelope and with condensed chromatin) or showed signs of proliferation, such as nuclei with an “8” shape or very elongated nuclei (Supplementary Fig. 2h). These results were also corroborated by the immunohistochemical analyses of phosphorylated histone H3, revealing proliferating cells in the EN (Supplementary Fig. 2p). Although it is known that endostyle cells proliferate [13], we found cSCs very close to, sometimes contacting, the endostyle cell basal lamina, but never delaminating from this organ, suggesting that cSCs homing this niche may come from the circulation, and do not derive from the endostyle itself.

By analysing histological serial sections of the endostyle and surrounding area in late secondary buds and primary buds, we retrieved cSCs in the body wall area surrounding the endostyle, and growing in number as buds progress in development.

Taken together, the above results clarify for the first time that the EN corresponds to a confined area located in the connective tissue ventral to the endostyle hosting resident cSCs which proliferate and differentiate.

## The body wall hosts cSCs aggregated in CIs, nodules and in the forming GN

CIs are agglomerates of cells, mostly phagocytes, located in the ventral mantle, lateral to the endostyle [14, 19]. cSCs were rare and scattered among phagocytes; some of them were in mitosis (Supplementary Fig. 2j–k). cSCs were also at the CIs periphery, in the form of nodules or scattered in the ventral body wall (Fig. 1d, Supplementary Fig. 2a–b, l-o). However, rows of differentiating cells, such as those seen in the EN, were never recognised, nor cells in differentiation. Histological serial sections of secondary and primary buds, revealed that CIs start to develop in primary buds, in the mid-cycle phase, as aggregates of phagocytes and a few cSCs, therefore earlier than previously reported [14, 19].

Moreover, we studied the morphology of the forming GN (Fig. 1g–h). When the early secondary bud is approaching the double vesicle stage [12], we could detect individual and groups of germ cells (both undifferentiated and some small previtellogenic oocytes) in the stalk sinus (called equatorial sinus within the secondary bud [18]) connecting the secondary bud to its parent, and in the lateral mantle of primary and secondary buds, the latter representing the GN. The study of serial sections evidenced that the secondary bud evaginates in an area very close to the gonad of the primary bud and that, before the stalk formation, the mantle is shared between the primary bud and its daughter bud. Some sections showed continuity of the germline between the parent gonadal region and its daughter bud. These results indicate that the GN receives the first homing germ cells directly from its parental gonad.

## Ampullae host cSCs and show a structure similar to the EN

To evaluate the presence of niches in the CCS, we serially cut a wide tunic area belonging to the same colony used to reconstruct the anatomy of the EN and CIs. We analysed 31 ampullae and the network of vessels connecting them to the marginal vessel and reconstructed their 3D structure (Fig. 2a–b; Supplementary Fig. 3a). We found that each ampulla contains a central sinus (often located toward the ventral ampulla wall), as direct continuation of the vessel from which the ampulla widens, whereas most of the ampulla volume is occupied by connective tissue (Fig. 2b–c). In this tissue, numerous hemocytes were recognizable (Fig. 2c–e): some of them were pigmented cells, many were hyaline amebocytes, some were differentiating hemocytes similar to those we found in the EN, most of them were cSCs. The presence of cSCs in connective tissue close to a sinus is a feature that ampullae share with the EN. A morphological comparison was carried out to assess the similarity among cSCs belonging to the different niches, finding that cSCs in the ampullae are comparable in nucleus/cytoplasm ratio and have similar histological features to those found in the EN and CI (Fig. 2g). We propose this microenvironment as the “ampulla niche” (AN). Most of cSCs were grouped in long bands in the ventral ampulla area, but in some cases organised around the sinus. Their reciprocal relationships (formation of rows, plasmalemma contacts) closely resemble those observed in cSCs in the EN. cSCs were found also close to the ampulla wall, together with hyaline amebocytes (Supplementary Fig. 3e). Some cSCs showed signs of proliferation, confirmed through immunohistochemistry as many cSCs were labelled by the anti-PH3 antibody (Fig. 2f) and at TEM. Single previtellogenic oocytes (sometimes degenerating or within phagocytic vacuoles in phagocytes; Fig. 2d–d’, Supplementary Fig. 3g) surrounded by primary follicle cells, and groups of 3-4 germ cells (Supplementary Fig. 3h) were also recognised, suggesting that both somatic and germ cSCs home in the AN. In live imaging videos (Supplementary Video 2), the presence of a sinus within each ampulla, with circulating cells, was clearly appreciable: only a limited volume of each ampulla was involved in the circulation, whereas most of the cells were scattered and immobile or slowly moving, making the ampullae similar in their structure to the EN and therefore credible as a niche where cSCs home. We finally evaluated the expression in ampullae of Myc, SoxB1, and Pou3, finding that cSCs are positive for the expression of these pluripotency and germ cell stemness markers (Fig. 2h–j), as seen in cSCs of the EN and CI in previous works [20].

We conclude that each ampulla has not only a role in hemolymph circulation, but is a SC niche containing both somatic and germ cSCs, showing morphological and molecular features similar to cSCs in the EN. Considering the large number of ampullae that a colony possesses and that they persist in the CCS independently from zooid life cycles, this niche represents a huge reservoir of cSCs.

## cSCs in the CCS participate in bud development

To characterise the participation of cSCs coming from the CCS in the development of newly forming individuals (primary and secondary buds), we designed a new method to exclusively label cells in the CCS (Fig. 3a, Supplementary Fig. 4a–c). This novel method involves the injection of a vital staining in the CCS of a colony in which all the zooids but one primary bud with its secondary buds were removed (called single-bud colony). The bud was also isolated from the CCS through the excision of the radial vessel joining it to the CCS. This method showed a high percentage (88%) of survival among isolated buds, which usually (93% of the surviving buds) reconnected to the marginal vessel within 24 hours thanks to the radial vessel regeneration, unless they were very far from the marginal vessel (Supplementary Fig. 4d–e). Injecting the vital staining CM-DiI (which is internalised in cells, hence avoiding leakage) into the vasculature before the radial vessel regeneration, allowed us to label only circulating hemocytes, including the cSCs. Therefore, labelled cells retrieved in the bud after the radial vessel regeneration exclusively came from the CCS.

**Fig. 3.**
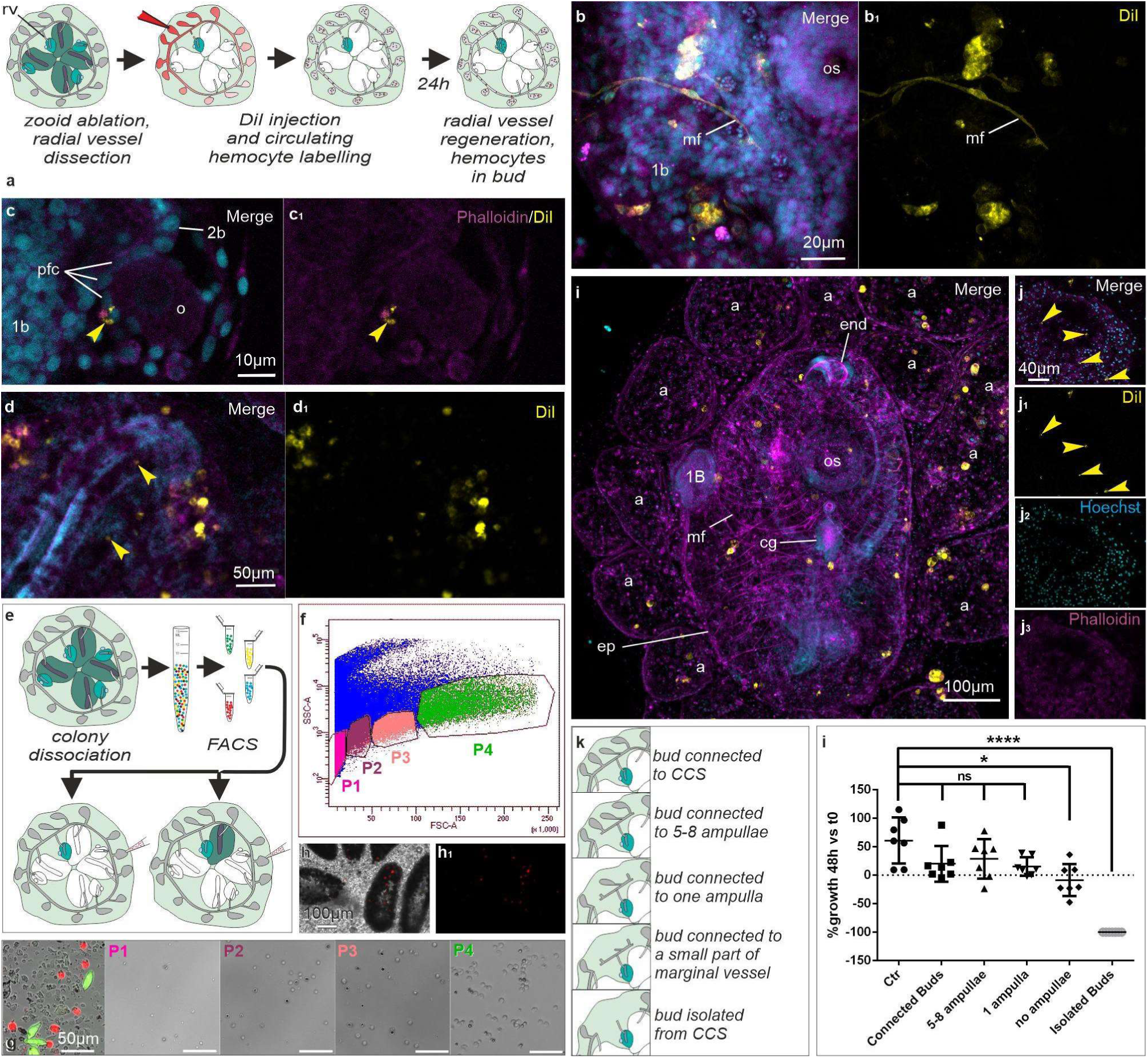
Contribution of circulating cSCs and ampullae in bud development. **a**. Illustration on the method to label *in vivo* circulating hemocytes. Ablated zooids are white. Only a primary bud (with its secondary bud) is left; its radial vessel (rd), connecting the bud to the colonial circulatory system, is sectioned but regenerates in 24 hours. See Supplementary Fig.1a for colony anatomy. **b,b_1_-d,d_1_.** Confocal images showing: a labelled muscle fibre (mf in b-b1) in a primary bud (1b), one day after reconnection (sample A_11); a labelled primary follicle cell (pfc; arrowheads in c-c1) in a primary bud (sample A_2); and labelled cells homing in the EN (arrowheads) (d-d1). Merge: Hoechst (blue), DiI (yellow), and Phalloidin (Magenta). o: oocyte, 2b: secondary bud. **e.** Schematic representation of the procedure followed for transplantation of sorted labelled cells. Cells were dissociated from a whole colony and sorted; the different cell populations were then injected into isogenic single-bud colonies (bottom left) and colonies with an adult zooid and its buds (bottom right). **f.** FACS gates. Single cells were selected, auto-fluorescent cells excluded, and cells with a low SSC profile chosen. **g.** Whole dissociated colony cells (right) and sorted populations P1-P4. Red and green cells: autofluorescent cells. Magnification is the same in all the panels. Labels as in (b). **h.** Ampullae (a) of a colony injected with FACS sorted cells homing some cSC (red spots, arrowheads). Live imaging. **i.** Adult zooid, injected with cells pertaining to P3, with several labelled cells (in yellow) in buds tissues and inside ampullae (a) (sample F621_P3). 1b: secondary bud. Frontal view, anterior at top, bud right side at left. end: endostyle; ep: epidermis; mf: muscle fiber; os: oral siphon. Confocal microscopy; colours as in b-b1. **j-j_3_.** Ampulla of a colony injected with FACS sorted cells homing some cSCs (yellow spots, arrowheads). Magnification is the same in j-j**_3_**. **k.** Experiments of bud isolation performed to study the contribution of the AN to bud survival and development. **i.** Isolated bud growth compared to the growth of control buds not isolated from the CCS. Even a single ampulla guarantees bud growth, whereas buds deprived of ampullae grow significantly less. Buds isolated from the CCS do not survive. Kruskal-Wallis test approximate P value = 0.0001. Dunn’s multiple comparison test: ns = not significative; * = p-value<0.05; ****= p-value <0.001.

We injected 17 single-bud colonies. Among these, only two buds could not reconnect to the marginal vessel. In live-imaging observations (n=8 colonies, Supplementary Video 3) performed in the following 5-6 days, we detected, among labelled hemocytes, cells with features of cSCs (about 5 µm in diameter) in several tissues. These cells were: i) circulating in the haemolymph, both inside the bud and in the CCS (including ampullae) (n=8 single-bud colonies), ii) immobile or with limited movements, in ampullae and in the developing EN and CI of primary buds (n=5), and iii) in tissues of primary and secondary buds (n=7). To determine the exact location of labelled cSCs, we then analysed the samples through confocal microscopy (Fig. 3b–d, Supplementary Fig. 4f–e, Supplementary Table 1), fixing colonies 5-6 days post-injection. At that time, most of the buds had opened their syphons, becoming adult individuals. Confocal images showed the labelling of various haemocyte types (i.e., giant phagocytes) coming from the CCS, including the small and round cSCs. We could also see labelled cSCs from the CCS infiltrated in the EN and CI (n=9 colonies) in primary buds (Fig. 3d), again evidencing that these niches develop in early primary buds, and do not pertain only to the adults. We found labelled cells in the branchial epithelium (n=2) (Supplementary Fig. 4f–f’’) and in epidermis (n=4) (Supplementary Fig. 4h–h’’) of secondary buds (confirmed checking nuclei alignment in the orthogonal projections), indicating that circulating cells had undergone a mesenchymal-epithelial transition. The contribution of cSCs to the differentiation of the epidermis was not previously demonstrated at this resolution in B. schlosseri. Moreover, we found labelled follicular cells in the GN of primary and secondary buds (n=3, Fig. 3c) and muscle fibres in the body wall musculature of a primary bud (n=1, Fig. 3b), confirming their development from mesenchymal cells [21, 22]. Finally, in a late secondary bud, we detected labelled cells in the differentiating dorsal tube epithelium (n=1), *i.e.* the rudiment of the bud cerebral ganglion and the associated neural gland [23], where labelled cells formed a small cluster with lower DiI fluorescence intensity, suggesting they derived by mitosis from circulating precursor cells(Supplementary Fig. 4f–f’’).

We also wondered whether the presence of adult individuals, that hold the already identified stem cell niches (EN, CI, GN), could influence the participation of cells from the CCS in bud development (Supplementary Fig. 4i–i’’, Supplementary Table 1). Labelled cells in bud tissues were retrieved in the same locations even when their parent was present, supporting the hypothesis that cells participating in bud development are also stored in the AN. Indeed, the above reported results confirm the presence, in the CCS, of cSCs or progenitor cells contributing to several tissues in bud development, which in turn could explain the massive regenerative abilities found in *B. schlosseri* and many other colonial ascidians [1–5, 24–27].

## Transplanted cSCs home in the ampulla niche and participate to bud development

Our results show that the CCS comprises a stem cell niche, the AN, and confirm the contribution of cSCs coming from the CCS to bud development. Therefore, to better characterise the function of cSCs from the CCS and their relationship with the AN, we isolated small and non-granular cells (*i.e.*, the cSC) through FACS and transplanted them into isogenic colonies after labelling them with CM-DiI (Fig. 3e). In a first round of experiments, populations of auto fluorescent cells were discarded, as they could represent algae and pigmented differentiated cells. We isolated non-granular cells by selecting cells with the lowest SSC (Side-Scatter) profile, and identified four different populations of small cells in the FSC (Forward-Scatter) profile (Fig. 3f): P1, represented the smallest entities, containing mostly cell debris and a few cells with a mean diameter of 5.1 µm; P2 and P3 were made up of cells with a mean diameter of 5.9 µm and 6.5 µm, respectively; P4 was the population showing the highest FSC profile, comprising cells of 6.8 µm in mean diameter, and with a maximum diameter of 9 µm (Fig. 3g). After labelling, we injected the cells in isogenic single-bud colonies, which were then monitored through live imaging and fixed after 3-7 days (N=4 biological replicates, n=21 injected single-bud colonies).

Only one sample injected with P1 cells presented labelled cells infiltrated into bud tissues (Supplementary Table 2): these were localised in the developing EN and CI niches, and in the epidermis of the primary bud. The other colonies injected with P1 cells underwent bud resorption (n=2) or did not have any labelled cell into buds (n=1). This could be due to the fact that this population was probably mainly composed of cell debris, or of possibly other cells/material that did not benefit colonies (such as bacteria that usually live outside the colony). Cells pertaining to P2, when reinjected in isogenic colonies (n=7), were found in CI (n=2), in epidermis (n=4) (Supplementary Fig. 5a, Supplementary Table 2) and in GN, namely in supposedly primary follicle cells of an oocyte in a secondary bud (n=1, Supplementary Fig. 5b). P3 cells were retrieved in many different tissues in the injected samples (n=5): other than the epidermis (n=3), EN and CI (n=4), we found labelled cells in the body wall musculature (n=1) (Supplementary Fig. 5c, Supplementary Table 2), and in the forming brain (n=2) of primary buds, in particular in the zone between the neural gland and the cerebral ganglion. Finally, cells pertaining to P4 were found differentiated in the epidermis (n=2 out of 5 injected colonies, Supplementary Fig. 5d) and in GN (n=1, Supplementary Table 2).

In a second round of experiments, we enriched our population of cSCs using the same gates on the SSC/FSC profile, but selecting only cells positive to the alkaline phosphatase activity [16]. In live imaging videos, many labelled cells were found immobile not only in the EN and CI, but also in the ampullae (Supplementary Video 4, Fig. 3h), while at the confocal microscopy many other cells were found in the ampullae (Fig. 3i–j) but also differentiated in the epidermis and gonads (both the somatic and germ lineages) (Supplementary Table 2). The presence of adult individuals did not influence the contribution of transplanted cells to bud development. The above results support that, in the AN, a population of cSCs is stored, which can be considered pluripotent as a whole as contributing to germ line development and also to several somatic lineages, such as muscle, epidermis and neurons.

## Ampullae are necessary for bud survival

To understand how important the contribution of the AN in development is, we generated single-bud colonies, varying the extension of CCS elements connected to the bud (Fig. 3k). We studied the fate of buds totally isolated from the colonial vasculature (isolated buds) (n=7), and buds that could take advantage of: a few (5-8) ampullae (5-8 ampullae buds) (n=7), only one ampulla (single ampulla buds) (n=5), a fragment of the marginal vessel without ampullae (no ampullae buds) (n=5), and the whole marginal vessel and ampullae (connected buds) (n=7). Colonies chosen for this experiment were in early-cycle and their primary buds had the heart beating. In all the experiments, colonies were not fed; anyway, buds had their siphons close, therefore could not feed.

Results highlight that, after 48 hours, buds connected to only a few or even just one ampulla were still growing, without showing any significant difference with the control buds in normal conditions, or with buds left attached to the whole CCS (Fig. 3l). Buds that were completely isolated from the vasculature were completely resorbed in 48 hours (Fig. 3l). To exclude the possibility that such an outcome was the result of the lack of nutrients stored in the circulatory system, we also compared the fate of buds left attached to the marginal vessel, but in which all the ampullae were removed. These buds stopped their growth after 48 hours and some of them started the reabsorption process, showing a significant difference in their growth when compared to the control group. Buds with only one ampulla however, did not reach the adult phase, as much as the “no ampullae-buds”, which however stopped growing earlier.

Buds connected to the whole CCS or to 5-8 ampullae were able to survive and develop normally, reaching the syphon opening stage (adulthood) with a little delay compared to the normal cycle timing, but without any significant difference both in terms of growth rate and days required for reaching adulthood. Conversely, isolated buds stopped heart beating soon after the surgical manipulation, and in 24-48 hours the internal structures were no longer recognizable. After 72 hours bud residuals were unrecognisable in all the observed cases, highlighting an important role of the haemolymph and its cells in the development of buds, and pointing out that when buds are connected to even a small number of ampullae they are able to survive and grow normally.

The above results highlight the influence of ampullae in supporting the development of individuals of the colony, since only a few ampullae are sufficient to sustain bud growth and even only one ampulla can initially sustain a growing phase, corroborating their role as SC niches.

## Discussion

*B. schlosseri,* as many other colonial tunicate species, show high regenerative capabilities culminating in whole body regeneration [1, 2, 5, 24–29]. This regenerative process happens in the circulatory system, when all the zooids are removed. However, SC niches in this species were identified only in the transitory structures of adults, fated to be resorbed [13, 14, 16]. In this work, we characterise the ampullae of the circulatory system as a new permanent SC niche, showing that cSCs which participate in bud development home in these structures, proliferate and differentiate. Moreover, by surgical manipulating the colonies, we proved that ampullae are an essential component for guaranteeing the growth of developing individuals. In the animal kingdom, various types of SC niches have been identified [30]. Many aquatic invertebrate species have no spatially confined microenvironments where SCs are stored and SCs fate is determined through intrinsic signals or through interactions with adjacent cells and signalling factors [31–33]. In this paper, we provide clear evidence that cSCs niches are present in *B. schlosseri* in the form of confined areas. Niches of *B. schlosseri* are therefore similar to the vertebrate counterparts, suggesting a similar way to maintain and store cSCs between *B. schlosseri* and vertebrates. Indeed, in a previous work [20] we characterised the expression, in cSCs in *B. schlosseri* EN and CI, of Yamanaka factors orthologues (*SoxB1*, *Pou3* and *Myc*), capable of reprogramming differentiated cells [34] and expressed constitutively by embryonic pluripotent SCs in mammals [35–37]. Rinkevich and collaborators [14] identified many vertebrate SC markers expressed by cSCs in the same niches. Although the lack of a precise anatomical characterization led the authors to assign cSCs, actually homing in the EN or circulating in the blood sinus, to the CIs, *in situ* hybridizations show the expression of genes such as *Piwi*, *Vasa* and *Stat* by cSCs in both EN and CI, suggesting that both these niches store cSCs. The similarities between *B. schlosseri* and mammals in the physical and molecular SC niches properties open to new comparative analyses on SCs maintenance in this organism. In this work we confirm that haemocytes participate in bud development even during the normal asexual cycle. Indeed, we proved that cells from the CCS infiltrate in primary and secondary bud territories, contributing to the development of several tissues, of both somatic (such as epidermis, muscle, egg envelopes, tunic cells) and germinal (e.g., oocytes) lineages, and in the EN, CI and GN. As recently shown [38], we confirmed the differentiation in the nervous system of cells coming from the CCS. Considering the developmental event occurring at the stage of cSCs transplantation and the retrieval stage, we conclude that many cSCs underwent mesenchymal to epithelial transition. The contribution to bud development occurred regardless of the presence of adult niches (EN, CI, and GN).

We demonstrated the presence of blood sinuses inside the ampullae evidencing that only a small area of each ampulla is involved in hemocyte circulation, whereas most of its volume is occupied by connective tissue, where cSCs are stored, some of them expressing *cMyc*, *Oct4* and *Sox2*. cSCs were found frequently in association with the ampulla wall and hyaline amoebocytes, possible components of the niche. The transplantation in isogenic colonies of cSCs isolated through FACS further proved that cells contributing to tissue development, also home inside the ampullae. Moreover, the experiments of bud isolation showed that the presence of ampullae influence the growth of isolated individuals, further proving their importance during development. Therefore, we show that ampullae are a SC niche, hosting both somatic and germ cSCs. The AN represents the first permanent niche described in colonial ascidians, since these structures store cSCs even during the resorption of adult individuals, when the EN, CI, and GN disappear from the old zooids and are replaced by the corresponding niches of the new maturing adult generation. Considering the number of ampullae per colony, and the huge amount of cSCs hosted by each ampulla, such niche in *B. schlosseri* and, likely, in closely related species (such as *Botrylloides sp.*), could be referred to as the niche supporting the extensive SC demand and use during the blastogenic development and whole-body regeneration, explaining the fundamental role of the CCS in development and regeneration.

The similarities between tunicate and vertebrate niches, coupled with the regenerative abilities shown by colonial tunicates and the close phylogenetic relationship between these simple chordates and vertebrates, highlight the importance of studying *B. schlosseri* and closely related species, as model systems for stem cell maintenance and regeneration.

## METHODS

### *B. schlosseri* colonies collection, rearing and cloning

Colonies of *B. schlosseri* were collected from the lagoon of Venice and reared in laboratory conditions at a constant temperature of 20°C. After collection, they were detached from their natural substrate (usually marine plants pertaining to the genus *Zoostera*, mussel shells or solitary sea squirts) with the aid of a razor blade and transferred to glass slides. They were kept in 10L aquaria, in which seawater was changed once a week. Colonies were feeded every day with microalgae (*Isochrysis galbana* and *Tetraselmis chuii*) cultured in the lab. To obtain isogenic colonies, large colonies were split by cutting the tunic and its circulatory system avoiding injuring the zooids. The resulting fragments were carefully detached from the glass with a razor blade, and transferred to separate glass supports. Colonies used for live imaging were left to adhere to microscopy plates (WillCo-dishes). Colonies used for surgical manipulations (isolation from vasculature or from other zooids) remained in filtered seawater (FSW) and were not fed until the bud reached adulthood and opened their syphons.

### Three-dimensional (3D) reconstructions

For the reconstruction of EN, CI and ampullae, a colony at the developmental phase 9/8/3 was used. Another colony at the same phase was used to reconstruct the GN in the secondary bud. Colonies, carefully selected as representative of the phase, were fixed for 24 h in a solution containing 1.7% glutaraldehyde and 1.7% sodium chloride in 0.2M sodium cacodylate buffer (pH 7.2). After two washes in sodium cacodylate buffer, the samples were post-fixed with osmium tetroxide solution and dehydrated through a graded ethanol series. After dehydration, samples were embedded in Epon resin (propylene oxide 1:1 for 1 h at 45°C and then twice with Epon 100% for 1 h). Samples were left to polymerize for three days at increasing temperatures (37°C-60°C) in silicone moulds, the excess of resin was removed, and samples were oriented. Sections 1 µm thick were obtained using an ultramicrotome (LKB Ultratome V) equipped with an Histo-Jumbo diamond knife (Diatome). Chains of sections were created through the application of glue at the base of the embedded sample, so as to attach sections one after another while cutting. Chains were collected and stuck to glass slides. Sections were then labelled with 1% Toluidine blue solution, photographed using a Leica 5000B optical microscope, and aligned using the software Adobe Photoshop CS. Using a graphic tablet, each individual organ was manually drawn on each section in the software Amira, which finally elaborated the final data, obtaining the 3D reconstruction of the sample. For the EN and CI 3D reconstruction, we performed 472 serial sections, to reconstruct part of the endostyle (corresponding to its medial-posterior portion, at level of the 6th-8th row of stigmata) and the surrounding organs. For the study of the ampullae and of the GN we performed 2720 and 75 sections, respectively. The diameter of cytoplasm and nucleus was measured for cSCs in the EN (13 cells), in CI (27 cells) and AN (13 cells), and the nucleus:cytoplasm ratio was calculated. For the analysis regarding differences in the nucleus:cytoplasm ratio as well as in the nucleus and cytoplasm diameter of haemoblasts from different compartments, a one-way ANOVA was applied after check of ANOVA assumptions.

### Transmission Electron Microscopy (TEM)

Colonies used for TEM were fixed and embedded in resin as described above. Ultrathin sections (80 nm thick) were then labelled with uranyl acetate and lead citrate. Photomicrographs were taken with a FEI Tecnai G12 electron microscope operating at 100 kV. Images were acquired with a Veleta (Olympus Soft Imaging System) digital camera.

### Isolation of buds from the vasculature

Colonies at developmental phases 9/8/2-3, consisting of only one or two systems and showing limited auto-fluorescence, were chosen. Before the surgical manipulation, the number of adult zooids and primary buds was annotated, as a reference for physiological colony conditions. Then, with the aid of 31G needle, all adult zooids and all the buds, except one primary bud with its budlets (usually represented by the right secondary bud, sometimes both the right and left secondary buds), were removed, creating a “single-bud colony”. In the experiments comprising the DiI injection in the vasculature, the remaining bud was also isolated from the colonial vasculature through the cutting of its radial vessel. We measured the distance between the isolated primary bud and its marginal vessels just after the surgery and every 12-24 hours. Statistical analyses were performed with the software GraphPad PRISM 6. We applied unpaired non-parametric Mann-Whitney U test to infer whether distance or physiological conditions influenced the reconnection of buds to the marginal vessel (Supplementary Fig. 4d–e). After 24 hours, the radial vessel regenerated in the majority of manipulated colonies, and the bud was again connected to the marginal vessel. In another set of experiments, a single adult and its buds were left, and the connections with the marginal vessel were destroyed through the cutting of the radial vessels of the adult and the primary buds.

### DiI injection and live imaging of single-bud colonies

In single-bud colonies, after 1-2 hours from the isolation of buds from the vasculature, about 5µl of CM-DiI dye (4.8 µM in FSW with 0.05% of phenol red) were injected in vascular ampullae, in this way labelling only circulating cells. Once the bud was reconnected to the marginal vessel, we started to monitor bud and budlets’ tissues to retrieve the presence of immobile labelled cells, which were therefore infiltrated from the circulation into tissues. Movies were taken using a Leica DMI4000B inverted microscope, with the microscopy plates filled with FSW and sealed with parafilm, to ensure the survival of colonies while being observed. Frames were taken with the smallest interval possible (ranging around 3 seconds) for 5 minutes, and combining brightfield images with epifluorescence acquisitions, using the filter cube N2.1 (Excitation: BP 515-560; Emission: LP 590). Once immobile cells could be observed in bud tissues, the colonies were fixed with 4% paraformaldehyde in 5% MOPS buffer solution (pH 7-7.5).

### Colony dissociation and FACS

Colonies used for the isolation of cSCs at FACS were dissociated as described by Laird (2005). Briefly, colonies were finely cut with a razor blade while immersed in *Botryllus* buffer (10mM Cystein, 25mM Hepes, 50mM EDTA in artificial sea water, pH 7.5). After filtration in 40 µm pore-size filters, cells were pelletted at 500g for 10 min and washed twice in freshly prepared 0.01M sodium citrate solution in artificial sea water. They were sorted in the same solution to prevent cell aggregation. In FACS experiments, we gated to exclude auto fluorescent cells in 555 and 488 channels, and only cells with a low SSC/FSC profile, as corresponding to small, non-granular cells were sorted. Four populations of cSCs were selected, as clearly separated in the SSC/FSC graph. In another series of experiments, colonies were labelled with propidium iodide (Invitrogen™ eBioscience™) to exclude dead cells and with Alkaline Phosphatase Live Stain (Thermo Fisher™) to select only the cells displaying a high activity of the alkaline phosphatase enzyme. As control, an unstained colony was used to design the gates and to avoid sorting autofluorescent cells. The same four populations with a low SSC/FSC profile were selected, but considering only positive cells to the alkaline phosphatase activity.

### Labelling and injection of cSCs

Sorted population of cSCs were pelleted and stained with CM-DiI dye (4.8 µM in artificial sea water), following the supplier protocol, and reinjected in isogenic single-bud colonies, in which all the buds and adult zooids were removed, except for a primary bud with its secondary buds, or except an adult with its buds, attached to the colonial vasculature. We injected 10.000 cells in the ampullae of each single-bud colony, using 0.05% phenol red in FSW to make the injected solution visible inside vessels and verify the success of the microinjection. We monitored the injected colonies through live imaging and fixed in 4% paraformaldehyde and 5% MOPS buffer solution (pH 7-7.5), when labelled cells were found immobile within the adult individual, the bud or its budlets.

### Labelling for confocal microscopy

After fixation, colonies were washed twice in phosphate-buffered saline (PBS: 1.37 M NaCl, 0.03 M KCl, 0.015 M KH_2_PO_4_, 0.065 M Na_2_HPO_4_, pH 7.2) and treated with Triton X 0.1% in PBS for 30 min. After two washes in PBS containing 0.1% Tween 20 (PBST), they were stained with Alexafluor-655 Phalloidin (ThermoFisher) 1:50 for 3 h, then washed again and stained with Hoechst for 30 minutes. Samples were mounted using Vectashield and images acquired with a Leica SP5 confocal inverted microscope or with a Zeiss LSM 700 confocal microscopy for upright acquisitions. Cells positive for DiI staining were annotated only when we were sure that they were infiltrated into the tissues (verified by orthogonal projections of the stacks and by the continuity and alignment with close cells and nuclei, using the software ImageJ). Phagocytes and pigmented cells, that could exhibit autofluorescence, were not considered in the results.

Colonies used for the identification of proliferating cells were fixed in 4% PFA in MOPS buffer (0.5 M MOPS, 2.5M NaCl, 5 mM MgSO4, 10 mM EGTA) overnight. To allow for better reagent penetration within tissues, after two washes in PBS, adult zooids were removed, and placed in 1.5 ml tubes. Adults and the remaining colony attached to the microscopy slide were treated separately. The initial steps of the protocol were the same listed above, with the addition of an overnight incubation with anti-phospho-histone H3 primary antibody (GTX128116, GeneTex, Irvine, CA, USA) after Triton X treatment. Secondary antibody staining (1:200) was performed together with the 3-hours phalloidin incubation. All acquired images were then elaborated using the software Fiji (Image J).

### *In situ* hybridisation

Whole mount *in situ* hybridizations were performed for *SoxB1, Pou3* and *Myc* using probes designed as previously reported (Vanni et al., 2022). Specimens were firstly fixed overnight using 4% paraformaldehyde in MOPS buffer (0.5M MOPS, 2.5M NaCl, 5mM MgSO4, 10mM EGTA), dehydrated using graded PBST/methanol solutions (50%, 70% and 100%) then dipped in xylene for 30 min to enhance the transparency of the samples and rehydrated with graded PBST/methanol (100%, 90%, 80%, 50%, 30%), with incubations of 15 min per step. Samples were then incubated for 30 min in 10 µg/ml Proteinase K in PBST at 37°C, washed three times in PBS and treated with 4% paraformaldehyde plus 0.2% glutaraldehyde in PBST for 30 min. They were then incubated for 1 h at 63°C while in the hybridisation mix (Amresco), and then overnight with digoxigenin (DIG) labelled riboprobes, at a concentration of 1μg/ml. They were then washed twice for 30 min in washing solution 1 (WS1: SSC (0.3 M NaCl, 40 mM sodium citrate, pH 4.5), SDS 10%, formamide 50%) at a temperature of 60°C, washed twice for 10 min in WS1 and washing solution 2 (WS2: 0.5M NaCl, 10 mM Tris-HCl, Tween-20 0.1%) at 60°C, twice in WS2 for 5 min at 37°C, then incubated for 30 min at 37°C in WS2 and RNase (final concentration 20µg/ml). After a short washing step of 5 min at 37°C with WS2, samples were again treated twice with WS1+WS2, once with WS1, once with WS1+Tris-buffered saline containing Tween (TBST: 137 mM NaCl, 2,7 mM KCl, 25 mM Tris-HCl, 0.1% Tween-20), twice with TBST for 10 min at 60°C for each washing step. Specimens were then treated three times with TBST for 5 min at room temperature (RT), incubated in 1% powdered milk in TBST (blocking solution) for 4 h at 4°C, and incubated for 12-16 h in the blocking solution and antidigoxigenin antibody conjugated with alkaline phosphatase (Ab-antiDIG) at 4°C. After a few washes in TBST, staining was performed in NTM (2M Tris-HCl, 2M NaCl, 2M MgCl2) and NBT-BCIP (Sigma, final concentration 0.3%), for 24 h at room temperature, while protected from light. After an incubation of 10 min in NTM at RT and in Tris-EDTA for 10 min, samples were washed twice (5 min at RT) dehydrated for the embedding procedure in graded ethanol in PBST (50%, 70%, 90% and 100%) and in xylene for 30 min at RT and 30 min 60°C. Finally, specimens were embedded in Paraplast Xtra (Sigma-Aldrich, St. Louis, MO, USA) and cut to 7 µm sections. Images were acquired using a Leica 5000B microscope. Control experiments were treated with sense probes.

### Experiments on isolated, connected or partially connected buds

For this experiment, five groups of single-bud colonies were treated differently, varying the amount of colonial vasculature they were connected to. In the first group of samples (defined as “connected buds”, n=7), the bud and the blood vessels were left intact. In the second group (defined as “5-8 ampullae buds”), two cuts in the marginal vessel were performed, leaving in this way the bud attached to a short tract of the vessel comprising a number of ampullae varying from 5 to 8. In the third group (defined as “single ampulla buds”) buds were left attached to a short fragment of the marginal vessel where all the ampullae were removed except one. In the fourth group (defined as “no ampullae buds”) buds were left attached to a fragment of the marginal vessel and all the ampullae were removed. In the fifth group (defined as “isolated buds”), the marginal vessel around the bud left in the colony was destroyed, so that the individual could not benefit from the support of colony hemolymph. In treated colonies, we checked daily the stage shift and the bud growth, measured considering the antero-posterior length of the primary bud, and comparing it to the bud length right before the surgery. Statistical analyses were performed using the software GraphPad PRISM 6. After checking that ANOVA assumptions were violated, we applied the Kruskall-Wallis test to compare the growth after 48 hours among buds which were surgically manipulated and control buds in intact colonies.

## Acknowledgments

We thank A. Cabrelle, N. Tecchio, S. Deppieri, S. Domenichi, C. Roncato, M. Bellio, F. Passini, G. Intorcia, for technical advices and data collection; the DiBio imaging facility (Department of Biology – University of Padova) for EM samples preparation; A. Spagnuolo, G. Fusco, L. Dalla Valle, B. Rosental, A. Voskoboynik, and J. Solana for helpful discussion.

## Funding

This research was supported by grants from the Italian MIUR (BIRD PRID 2021) to L.M. and COST Action MARISTEM 16203 to L.M. and L.B.

## Contributions

V.V. and L.M. conceived and planned the project with inputs from G.M.. F.C. and L.M. performed 3D reconstruction and TEM. V.V. performed surgical manipulation of *B. schlosseri* colonies, confocal microscopy and *in situ* hybridization. V.V. and A.P. performed FACS sorting and live imaging. D.A., F.G. and V.V. performed statistical analyses. V.V., L.M. and G.M. wrote the manuscript. All the authors approved the final manuscript.

**Supplementary Fig. 1.**
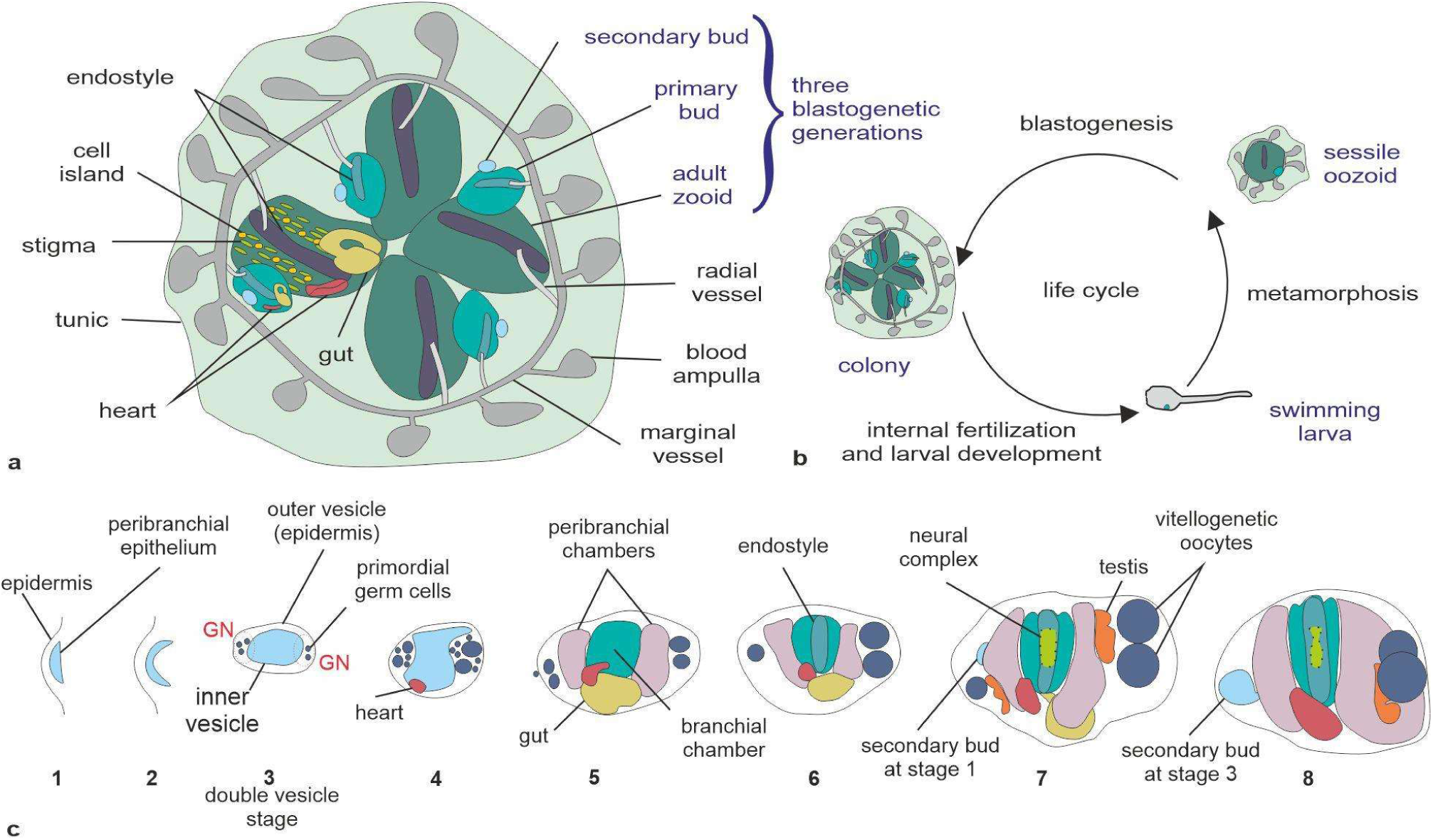
*B. schlosseri* anatomy and life cycle. **a**. Schematic representation of a *B. schlosseri* colony, composed of a single system. A system consists of a star-shaped group of adult zooids (four in this case), sharing the cloacal opening at the centre of the system. A colony can include hundreds of systems. The three generations composing the colony are highlighted in blue. **b.** Life cycle of *B. schlosseri*. A tadpole larva, possessing the typical chordate body plan, is generated by internal fertilisation. After a few hours of free swimming, the larva adheres to a substrate and metamorphoses into a sessile, filter-feeding individual, the oozooid, the founder of the new colony, which exhibits a new individual (blastozooid in form of primary bud, light green) produced through asexual reproduction (blastogenesis) (also recognizable as secondary bud in the larva). **c.** Illustration of bud development in ventral view. 1-6: developmental stages of secondary bud; 7-8: developmental stages of primary bud. The neural complex (in light green) in the primary bud is dotted, being located dorsally to the brachial chamber. GN in secondary bud stage 3: gonadal niche. Modified from Vanni et al., 2022.

**Supplementary Fig. 2.**
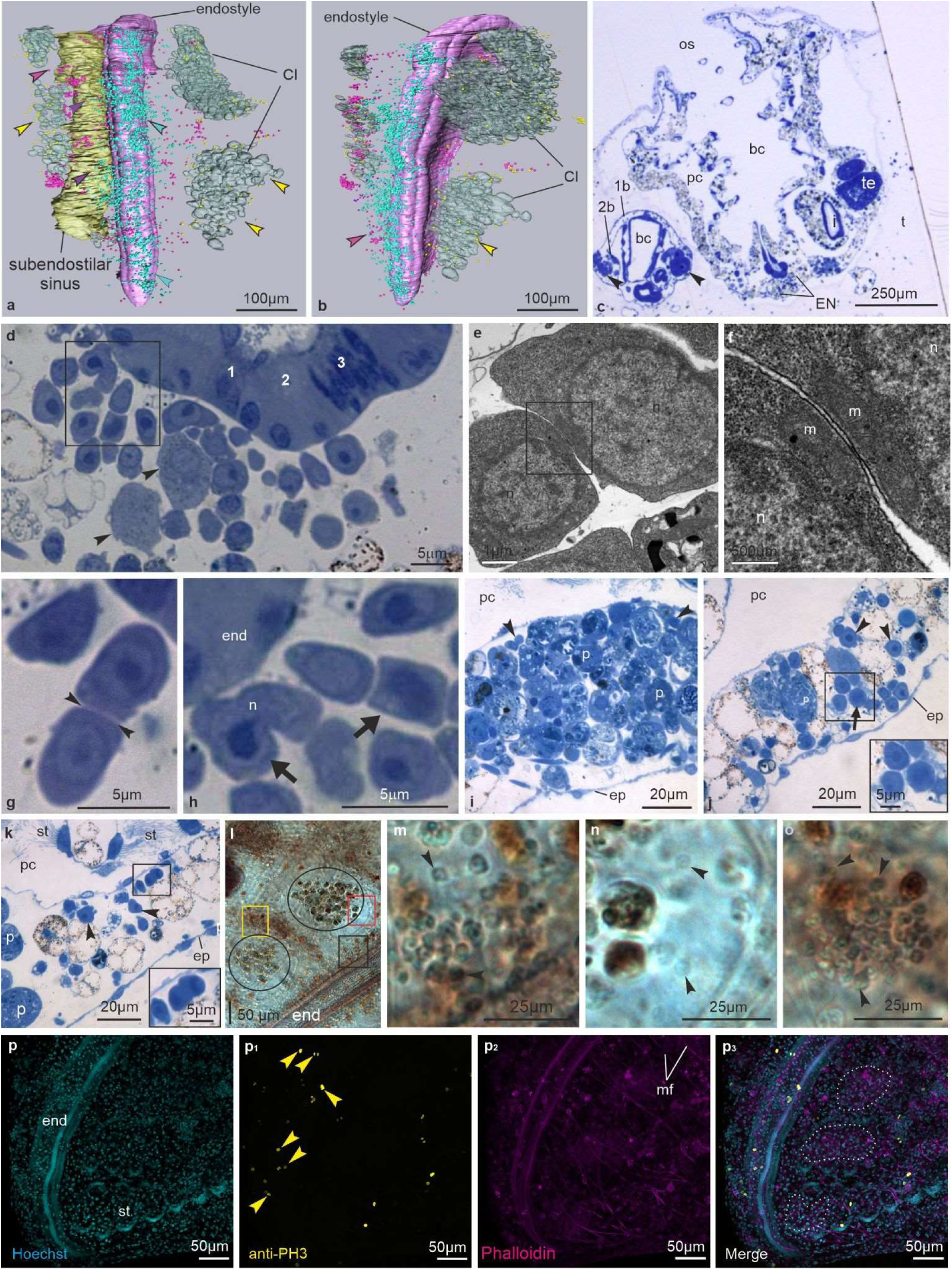
EN and CI: morphology and cell proliferation. **a-b.** 3D reconstruction of the posterior half of the EN and the surrounding tissues of an adult zooid (colony phase 9/8/3). **a**: the epidermis, the heart, the branchial and peribranchial chambers, and the ciliated epithelium of stigmata were omitted with respect to Fig. 1d; ventral view. In (**b**), also the subendostilar sinus was omitted; lateral view. Note that the EN is rich in cSCs (blue spots, blue arrowheads). A few cSCs are within the sinus (purple spots, purple arrowheads). Some cSCs are in the connective tissue (fuchsia spots, fuchsia arrowhead), also aggregated in small nodules (black arrowheads). A few cSC are in CI (yellow spots, yellow arrowheads). **c.** Transverse histological section of an adult zooid with its primary (1b) and secondary (2b) buds belonging to the individual used for the 3D reconstruction. In the primary bud, note the bilateral GN (arrowheads), more developed on the left body wall than the right one (the right side is marked by the secondary bud, 2b). Toluidine blue. bc: branchial chamber; EN: endostyle niche; i: intestine; os: oral syphon; pc: peribranchial chamber; t: tunic; te: testis lobules. **d**. Details of EN. Some cells (arrowheads) show signs of differentiation, such as granules in cytoplasm. Squared area corresponds to that one shown in (**h**). Toluidine blue. **e-f.** Two cSCs in the EN, with a large nucleus (n) encircled by a thin area of homogeneous cytoplasm, rich in free ribosomes. The square area in (**e**) is enlarged in (**f**) to show the small contact between the two cSC. Fibrillar material is recognizable in the cleft, and mitochondria (m) are very close to the cell membranes. TEM. **g**. Pair of cSCs in the EN strictly in contact by means of their plasma membrane (arrowheads). **h.** Two mitotic cSCs in the EN (arrows): the left one has an elongated nucleus (n) in a figure of eight, the right one does not show a nuclear envelope and has condensed chromatin. Toluidine blue. **i-k**. Transverse sections of CI in the adult zooid shown in (**a**). **i**: CI with large phagocytes (p) and a few cSCs (arrowheads). **j**: peripheral view of a CI. Inset: a mitotic cSC (arrow) as not exhibiting a nuclear envelope. **k**: cSCs (enlarged in inset) in mitosis in the connective tissue close to a CI. Toluidine blue. ep: epidermis; pc: peribranchial chamber; st: stigma. **l-o.** Detail of an adult zooid to show the EN and CI (circles). Black square area is enlarged in (**m**), red square area in (**n**), yellow square area in (**o**), to show cSC in EN, in CI, and in a nodule in ventral body wall, respectively. Ventral view, whole mount colony, phase 9/8/5. **p-p_3_**. Anterior ventral view of a zooid treated with antibody anti-phosphorylated histone H3. Proliferating cells (yellow spots) are in the EN (arrowheads) and close to CI. Dotted lines in (**p_3_**): CI.

**Supplementary Fig. 3.**
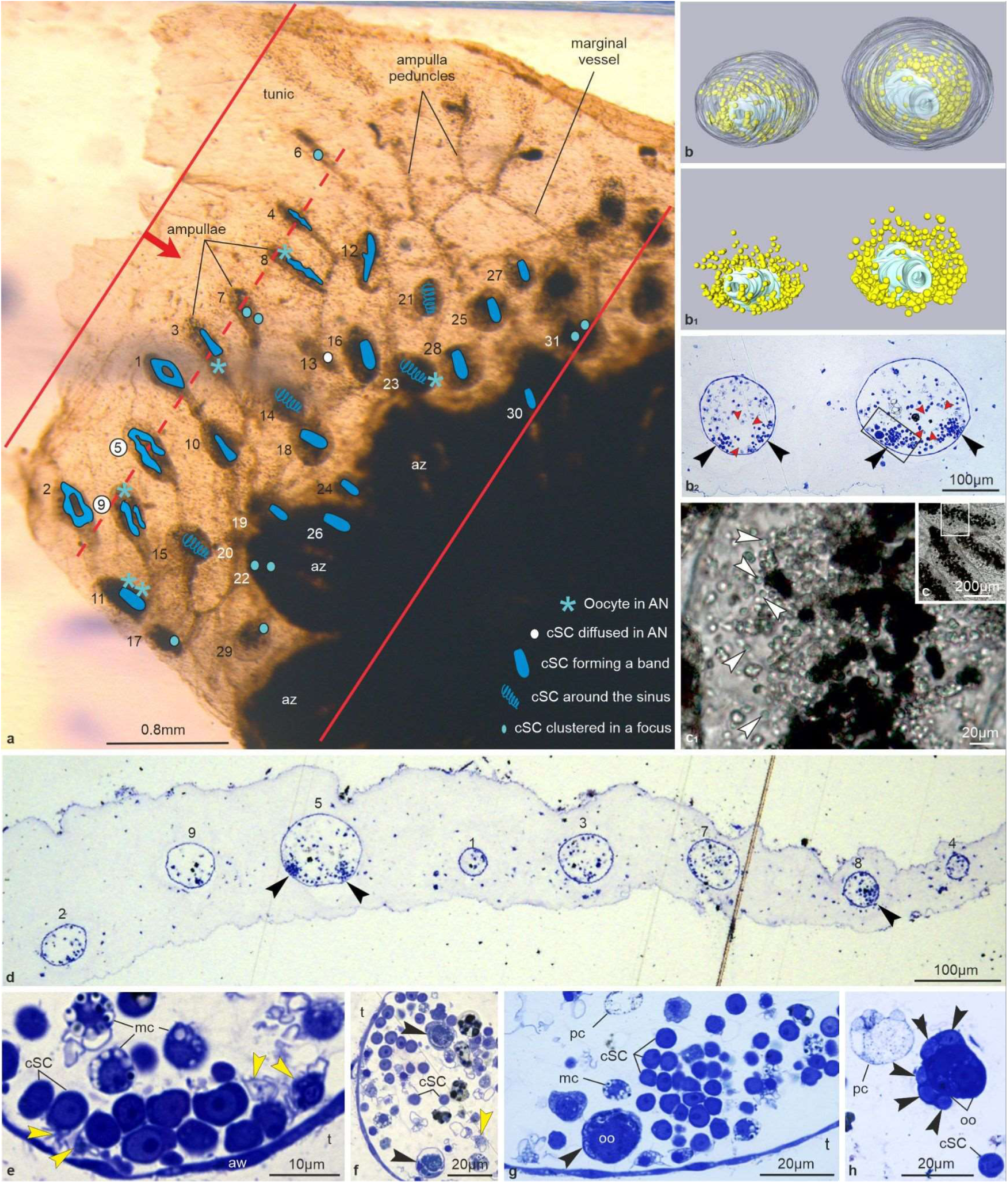
AN structure. **a.** Detail of the colony (ventral view) used for the analyses of the CCS, embedded in the resin block before sectioning. The tunic area included between the two red lines was serially sectioned, producing 2720 sections 1μm thick. The red arrow indicates the cutting direction; the dotted red line indicates the level of section shown in (**d**). 1-31 are the studied ampullae (numbered according to their appearance in serial sections); ampullae n. 9 and n. 5 have been 3D reconstructed; ampullae n. 22, 26, and 30 are hidden by adult zooids (az). Each ampulla contained a niche, whose shape has been approximately illustrated in blue (see legend at button right). Previtellogenic oocytes were recognized in ampullae n. 2, 8, 9, 11, and 23. **b-b_2_**. Ampullae n. 9 (left) and n. 5 (right) in 3D reconstructions from their apex in (**b**-**b_1_**), transverse section in (**b_2_**). The epidermis is transparent in (**b**) and has been omitted in (**b_1_**), to show the cSC (yellow spots). In (**b_2_**), the two sinuses in ampullae are indicated by red arrowheads, whereas clusters of cSC by black arrowheads; the squared area is enlarged in (**g**). Scale bar is the same in (**b**-**b_2_**). Toluidine blue in (**b_2_**). **c-c_1_**. Ampullae *in vivo* (from Fig. 2a and Supplementary Video 2). The squared area in (**c**) is enlarged in (**c_1_**) to show cSCs (white arrowheads) whose diameter is about 5 μm. Black spots: pigmented cells above the ampulla sinus. **d.** Section selected from the pool of 2720 sections used to study the CCS, cut at level of the dotted red line in (**a**). It shows 8 ampullae, numbered as in (**a**). Dorsal side at top. Clusters of cSCs (arrowheads) are mostly close to the ventral ampulla wall. **e-h.** Detail of AN belonging to different ampullae. In (**e**), note the relationships between cSCs and the ampulla wall (aw). Hyaline amebocytes (yellow arrowheads) are close to cSCs. (**f**): two large phagocytes (black arrowheads) close to a cluster of cSCs. Yellow arrowhead: hyaline amoebocyte. (**g**): enlargement of the square area shown in (**b_2_**). A degenerating previtellogenic oocyte (oo) surrounded by primary follicle cells (arrowhead) is close to cSCs. (**h**): a cluster of germ cells (arrowheads) and two oocytes (oo) in an AN. mc: differentiating morula cell; pc: pigmented cell; t: tunic.

**Supplementary Fig. 4.**
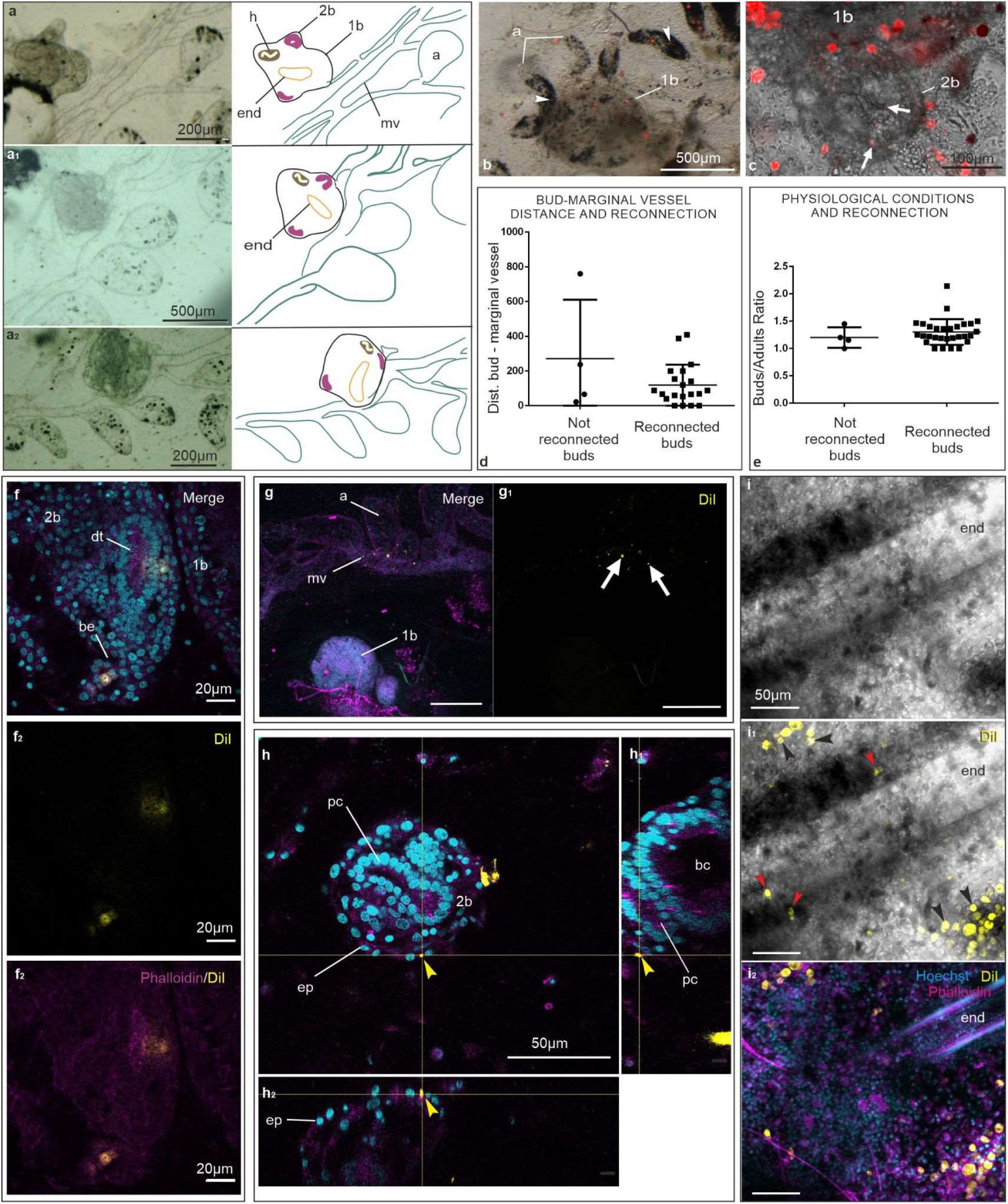
Dynamics of bud reconnection and participation of circulating cSCs to blastogenesis. **a-a_2_.** *In vivo* images and corresponding schematic illustrations of isolated primary bud (1b) reconnection to the CCS. After 24h from the experimental intervention, labelled hemocytes can invade the buds. Images taken just after the isolation of the bud (**a**), after 16 h (**a_1_**) and 24h (**a_2_**). 2b: secondary bud; a: ampulla; end: endostyle; h: heart; mv: marginal vessel. **b-c**. Primary (1b) and secondary (2b) isolated buds 72 hours after surgery, showing labelled cells (red spots) inside tissues. Stationary cSCs are in buds (arrows) and ampullae (a, arrowheads). Live imaging at stereomicroscope (**b**, sample A_20) and at inverted microscope (**c**, sample L_14). **d.** Dot plot showing the distance between buds and the marginal vessels in colonies that successfully or unsuccessfully reconnected to the CCS. The t-test did not show a significant difference between the two groups. **e.** Dot plot showing the number of buds per adult individual (used as a characteristic to assess the physiological condition of the colony) in buds that successfully or unsuccessfully reconnected to the CCS. The t-test did not show a significant difference between the two groups. **f-f_2_**. Confocal image showing labelled cells in the branchial epithelium (be) and dorsal tube (dt). Note in dorsal tube-labelled cells the dilution of the dye suggesting an occurred cell proliferation (sample L_14). Merge: DiI (yellow); Hoechst (blue); phalloidin (magenta). **g-g_1_**. Confocal image showing a bud (1b) that did not reconnect to the marginal vessel (mv) and was resorbing. Labelled cells were not retrieved inside bud tissues, while they were recognizable in the marginal vessel (mv; arrows in B’) (sample A_17). a: ampulla. Merge: DiI (yellow); Hoechst (blue); phalloidin (magenta). **h-h_2_**. Confocal image showing a labelled cell (arrowhead) in the epidermis (ep) of a secondary bud (2b) (sample A_6). **h_1_-h_2_**: orthogonal planes passing through the labelled epidermal cell, showing the alignment of the labelled cell nucleus with those of the adjacent epidermal cells, indicating labelled cell infiltration in the epithelium. DiI (yellow); Hoechst (blue); phalloidin (magenta). **i-i_2_**. Confocal microscopy images of labelled hemocytes located in the EN (red arrowheads) and CI (black arrowheads) of an adult zooid. Bright field on **i**, merge of bright field and DiI (yellow) in **i_1_**, merge of DiI (yellow), Hoechst (cyan) and phalloidin (magenta) on **i_2_**.

**Supplementary Table 1.** Synthesis of results obtained injecting DiI in ampullae of single-bud colonies. In the columns “Phase of injection” and “Phase of fixation” numbers in grey indicate the absence of the adult individuals in the colony. «Fixation day» indicates the number of days from the injection day. Territories in which labelled cells were found in adult zooids (az), primary buds (1b) and secondary buds (2b) are shown. In “Sample ID” column, L_ samples indicate colonies followed every day by live imaging; Ad_ samples indicate colonies in which adults were left, and were isolated from the marginal vessel.

| Sample ID | Phase of injection | Fixation day | Phase at fixation | Territories of adult zooids (az), primary buds (1b) and secondary buds (2b) in which labelled cells were found |  |  |  |  |  |
| --- | --- | --- | --- | --- | --- | --- | --- | --- | --- |
|  |  |  |  | EN + CI | Branchial Epithelium | Dorsal tube epithelium | Epidermis | Muscle fibres | GN |
| A 6 | 9/8/3 | 3 | 9/8/5 | 1b | 1b |  | 1b + 2b |  |  |
| A_10 | 9/8/3 | 2 | 9/8/3 | 1b |  |  |  |  |  |
| A_11 | 9/8/3 | 2 | 9/8/3 | 1b |  |  |  | 1b |  |
| A_12 | 9/8/2 | 3 | 8 (buds resorbed) | 1b |  |  |  |  |  |
| A_14 | 9/8/2 | 4 | 11/8/6 | 1b |  |  |  |  |  |
| A_16 | 9/8/2 | 4 | Not reconnected |  |  |  |  |  |  |
| A 17 | 9/8/2 |  | Not reconnected |  |  |  |  |  |  |
| A_19 | 9/8/3 | 4 | 11/8/6 | 1b |  |  |  |  |  |
| A 20 | 9/8/2 | 7 | 9/7/1 | az |  |  | 2b |  |  |
| L 2 | 9/8/2 | 7 | 9/7/1 | az |  |  |  |  | 1b |
| L 6 | 9/8/2 | 7 | 9/8/5 | 1b |  |  |  |  |  |
| L 9 | 9/8/3 | 8 | 11/8/6 |  |  |  |  |  | 2b |
| L 10 | 9/8/2 | 7 | 9/8/3 | 1b |  |  |  |  |  |
| L_14 | 9/8/3 | 3 | 11/8/6 | 1b | 2b | 2b |  |  | 2b |
| L_20 | 9/8/3 | 7 | 11/8/6 | 1b |  |  |  |  |  |
| L_24 | 9/8/2 | 15 | 9/7/1 |  |  |  |  |  |  |
| L_25 | 9/8/2 | 6 | 9/7/1 | az |  |  |  |  |  |
| Ad_1 | 9/7/1 | 5 | 9/8/2 | az |  |  | 1b |  |  |
| Ad 2 | 9/8/2 | 5 | 9/7/1 | az |  |  |  | 1b |  |
| Ad_3 | 9/8/2 | 8 | 9/8/4 | az |  |  |  |  |  |
| Ad 4 | 9/8/2 | 5 | 9/7/1 | az |  |  | az |  |  |
| Ad 5 | 9/8/2 | 4 | 9/8/2 | az |  |  |  |  |  |

**Supplementary Table 2.** Synthesis of results obtained injecting FACS sorted cells in ampullae of single-bud colonies. Sorted cells were small, non-granular and non-auto fluorescent, or with high levels of alkaline phosphatase activity. They were transplanted into isogenic colonies after labelling with CM-DiI. In the columns “Stage of Injection” and “Stage of fixation”, numbers in grey indicate the absence of the adult individuals in the colony. “Fixation day” indicates the number of days from the injection day. Territories in which labelled cells were found in adult zooids (az), primary buds (1b) and secondary buds (2b) are shown. In “colony ID” sample names starting with A indicate colonies in which adults were left, and P1, P2, P3, P4 indicate the sorted population injected in the colony. NA – not assessed.

| Colony ID | Injected Population | Stage of Injection | Fixation Day | Stage at Fixation | Territories of adult zooids (AD), primary buds (1B) and secondary buds (2B) in which labelled cells were found |  |  |  |  |  |  |
| --- | --- | --- | --- | --- | --- | --- | --- | --- | --- | --- | --- |
|  |  |  |  |  | EN | CI | Neural complex | Epidermis | Muscle fibers | GN | Immobile cells in AN? |
| F541_P1 | P1 | 9/8/2 | 3 | Resorbed |  |  |  |  |  |  | NA |
| F541_P2 | P2 | 9/8/4 | 4 | Resorbed |  |  |  |  |  |  | NA |
| F541_P3 | P3 | 9/8/4 | 4 | 9/8/2 |  |  |  |  |  |  | NA |
| F541_P4 | P4 | 9/8/2 | 4 | 9/8/5 |  |  | 1b |  |  |  | NA |
| F574_P1 | P1 | 9/8/5 | 5 | 9/8/3 | az | az |  | az |  |  | NA |
| F574_P2_A | P2 | 9/8/5 | 4 | 9/7/1 |  |  |  | 1b |  |  | NA |
| F574_P2_B | P2 | 9/8/3 | 5 | 9/7/1 | az | az |  |  |  | 1b | Y |
| F574_P2_C | P2 | 9/7/1 | 5 | 9/8/5 |  | 1b | 1b | 2b |  |  | NA |
| F574_P3 | P3 | 9/8/3 | 5 | 9/7/1 |  |  |  | 1b |  |  | NA |
| F574_P4 | P4 | 9/8/2 | 4 | 9/8/5 |  |  |  | 1b + 2b |  | 2b | NA |
| F621_P1 | P1 | 9/8/2 | 4 | 9/8/5 |  |  |  |  |  |  | NA |
| F621_P2_A | P2 | 9/8/4 | 4 | 9/7/1 |  |  |  | 1b |  |  | NA |
| F621_P3 | P3 | 9/8/4 | 4 | 9/8/2 |  |  |  | az | az |  | NA |
| F621_P4 | P4 | 9/8/2 | 4 | Resorbed |  |  |  |  |  |  | NA |
| ALCALINE FOSFATASE ACIVITY POSITIVE CELL POPULATIONS |  |  |  |  |  |  |  |  |  |  |  |
| FS1_P1 | P1 | 9/7/1 | 5 | 9/8/4 |  |  |  |  |  |  |  |
| AFS1_P1 | P1 | 9/7/1 | 5 | 9/8/4 |  |  |  |  |  |  |  |
| FS1_P2 | P2 | 11/8/6 | 5 | 9/8/3 |  |  |  |  |  |  | Y |
| AFS1_P2 | P2 | 9/8/2 | 5 | 9/8/5 | az |  |  |  |  |  |  |
| FS1_P3 | P3 | 9/8/2 | 5 | 9/8/5 |  |  |  |  |  | 1b | Y |
| AFS1_P3 | P3 | 9/8/2 | 5 | 11/8/6 | az | az |  |  |  |  | Y |
| FS1_P4 | P4 | 9/8/3 | 5 | 9/7/1 |  | az |  | 1b |  |  | Y |
| AFS1_P4 | P4 | 9/8/2 | 5 | 9/8/5 |  |  |  |  |  |  | Y |
| FS2_P1 | P1 | 9/8/5 | 6 | 9/8/3 |  |  |  |  |  |  |  |
| AFS2_P1 | P1 | 9/8/5 | 6 | 9/8/4 |  |  |  |  |  |  |  |
| FS2_P2 | P2 | 9/8/5 | 6 | 9/8/3 |  |  |  |  |  |  |  |
| AFS2_P2 | P2 | 9/8/5 | 6 | 9/8/3 | az |  |  |  |  |  |  |
| FS2_P3 | P3 | 9/8/4 | 6 | 9/8/2 | az |  |  |  |  |  |  |
| AFS2_P3 | P3 | 9/8/4 | 6 | 9/8/3 |  | az |  |  |  |  | Y |
| FS2_P4 | P4 | 9/8/4 | 6 | 9/8/2 |  |  |  |  |  | 1b | y |
| AFS2_P4 | P4 | 9/8/4 | 6 | 9/8/2 |  |  |  |  |  |  |  |
| FS3_P1 | P1 | 9/7/1 | 6 | 9/8/5 |  |  |  |  |  |  |  |
| AFS3_P1 | P1 | 9/7/1 | 6 | 9/8/5 |  | az |  |  |  |  |  |
| FS3_P2 | P2 | 9/8/3 | 6 | 9/7/1 |  | az |  |  |  |  |  |
| AFS3_P2 | P2 | 9/8/3 | 6 | 9/7/1 |  | az |  |  |  |  | Y |
| FS3_P3 | P3 | 11/8/6 | 6 | 9/8/4 | az | az |  |  |  |  | Y |
| AFS3_P3 | P3 | 11/8/6 | 6 | 9/8/4 | az+1b | az |  | az |  |  | Y |
| FS3_P4 | P4 | 9/7/1 | 6 | 9/8/4 | 1b |  |  | 1b |  |  |  |
| AFS3_P4 | P4 | 9/7/1 | 6 | 9/7/1 |  | az |  | 1b |  |  |  |

**Supplementary Fig. 5.**
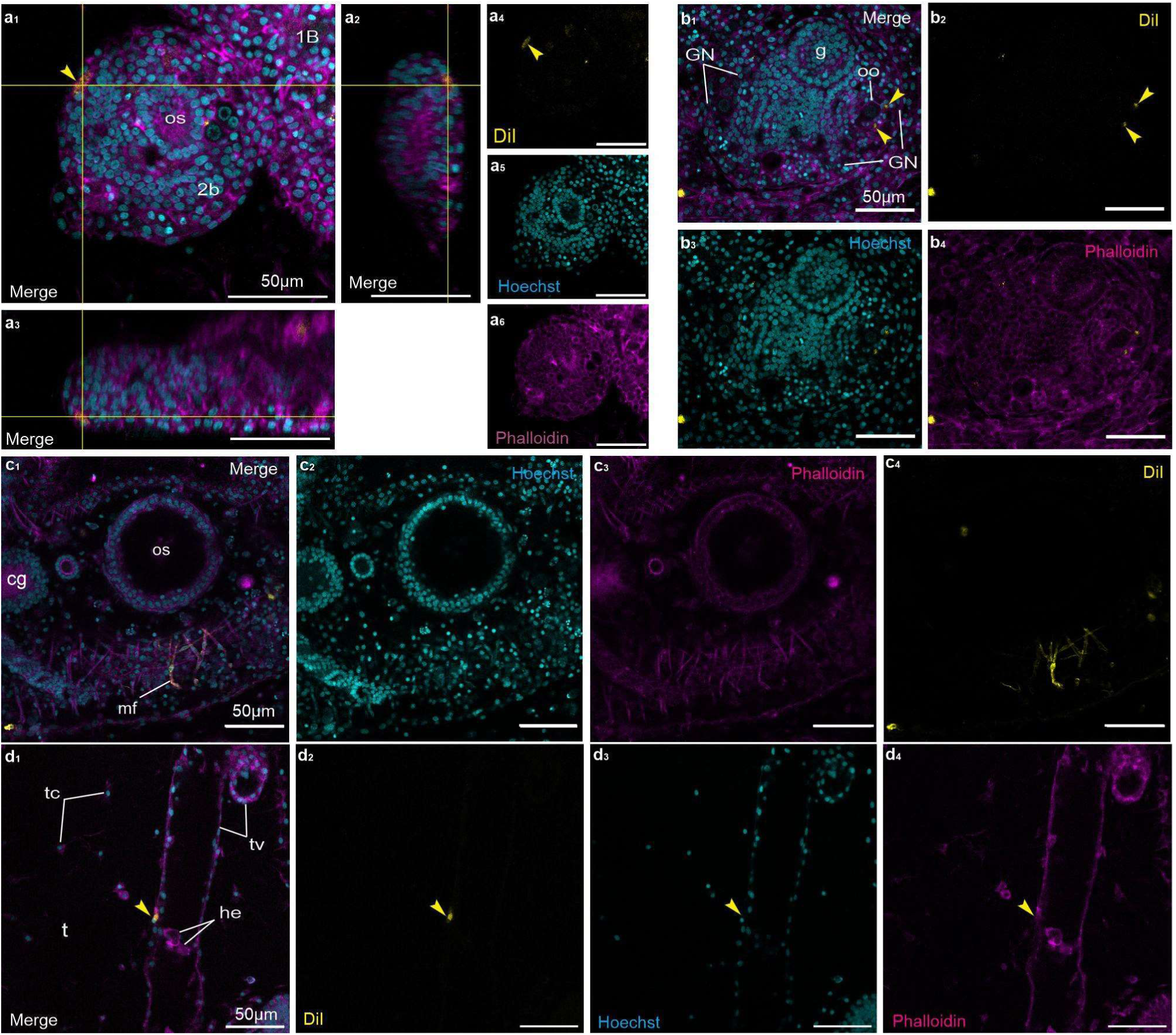
Participation of sorted cSC populations to bud development. **a**_1-6_ Sample F574_P2_C. Confocal microscopy. **a_1_**. Labelled epidermal cell (arrowhead) in a secondary bud (2b, stage 5), injected with cells pertaining to P2. The epithelial location of the labelled cell is evidenced also by the orthogonal projections in **a_2_** and **a_3_**. Frontal view, anterior at top-left. Merge: Hoechst (blue), DiI (yellow), and phalloidin (magenta). **a_4_-a_6_**. DiI (yellow), Hoescht (blue), and Phalloidin (magenta), respectively. Scale bar is the same in a_1_-a_6_. 1b: primary bud; 2b: secondary bud; os: oral siphon in secondary bud. **b**_1-4_. Secondary bud at GN level, injected with cells pertaining to P4. The GN is bilateral. (Sample F574_P4; frontal view, anterior at bottom-left, bud left side at right). Two supposedly primary follicle cells (arrowheads) belonging to a previtellogenetic oocyte (oo) are labelled. Confocal microscopy. g: gut. Colours as in B; scale bar is the same in b_1-4_. **c**_1-4_. A labelled muscle fibre (mf) in an adult zooid (sample F621_P3; frontal view, anterior at top right, bud right side at 8/2). cg: cerebral ganglion; os: oral siphon. Colours as in **a**. Scale bar is the same in c_1-4_. **d1-4.** Vascular epithelial cell labelled in a colony injected with P3 cells. Colours as in **a**. Scale bar is the same in all panels.

## SUPPLEMENTARY VIDEO LEGENDS

**Supplementary Video 1. Relationship between cSCs in EN and CI and the circulatory system.** Live imaging of an adult zooid in its ventral side. cSCs in EN and CI show limited movements and are not part of the subendostylar sinus, where cells can be seen moving very quickly, proving that cSCs home in the niches. Scale bar 50 µm.

**Supplementary Video 2. Ampullae are composed of a blood sinus surrounded by hemocytes.** Video showing the circulation inside the ampullae in ventral view. In each ampulla, a central sinus is surrounded by pigmented cells. cSCs, together with different types of hemocytes, are found around the sinus and show limited movement in comparison with hemocytes inside sinuses.

**Supplementary Video 3. Circulating cells infiltrate in bud tissues after DiI injection.** Live imaging time lapses of a single-bud colony injected with DiI and observed for 6 days. cSCs can be observed in the EN an CI as well as circulating in the hemolymph. Images were taken with the smallest interval possible (ranging around 3 seconds) for 5 minutes. Scale bar: 50 µm

**Supplementary Video 4. Sorted cSCs home in ampullae.** Live imaging video of a colony in which sorted and labelled cSCs contributing to bud development were transplanted. Immobile small cells can be seen inside ampullae, suggesting that they home in these structures. Scale bar: 100 µm

## Notes

### Competing Interest Statement

The authors have declared no competing interest.

